# General environmental heterogeneity as the explanation of sexuality? Comparative study shows that ancient asexual taxa are associated with both biotically and abiotically homogeneous environments

**DOI:** 10.1101/141317

**Authors:** Jan Toman, Jaroslav Flegr

**Affiliations:** Laboratory of Evolutionary Biology, Department of Philosophy and History of Sciences, Faculty of Science, Charles University in Prague, Vinicna 7, 128 00 Prague 2, Czech Republic

**Keywords:** ancient asexuals, asexual reproduction, Frozen evolution theory, habitat heterogeneity, sexual reproduction

## Abstract

Ecological theories of sexual reproduction assume that sexuality is advantageous in certain conditions, for example, in biotically or abiotically more heterogeneous environments. Such theories thus could be tested by comparative studies. However, the published results of these studies are rather unconvincing. Here we present the results of a new comparative study based solely on the ancient asexual clades. The association with biotically or abiotically homogeneous environments in these asexual clades was compared with the same association in their sister, or closely related, sexual clades. Using the conservative definition of ancient asexuals (i.e. age > 1 million years), we found six pairs for which relevant ecological data are available. The difference between the homogeneity type of environment associated with the sexual and asexual species was then compared in an exact binomial test. Based on available literature, the results showed that the vast majority of ancient asexual clades tend to be associated with biotically or abiotically, biotically, and abiotically more homogeneous environments than their sexual controls. In the exploratory part of the study, we found that the ancient asexuals often have durable resting stages, enabling life in subjectively homogeneous environments, live in the absence of intense biotic interactions, and are very often sedentary, inhabiting benthos and soil. The consequences of these findings for the ecological theories of sexual reproduction are discussed.

## Introduction

### Paradox of sexual reproduction

The term “sexual” may be assigned to a broad array of processes (see e.g. Redfield, 2001; Birky, 2009; Gorelick & Carpinone, 2009; Schön et al., 2009a). On the other hand, sex in its strict sense (i.e. amphimixis) is defined as the alternation of meiosis and syngamy (Butlin et al., 1998a).

Sexual reproduction (*sensu* amphimixis) is one of the most enigmatic phenomena in evolutionary biology, especially if its overwhelming predominance in eukaryotes is taken into account (see e.g. Williams, 1975; Maynard Smith, 1978; Bell, 1982; Meirmans & Strand, 2010). The reason is that it brings many obvious disadvantages in comparison to asexual reproduction— the well-known two-fold cost of sex being only the first and most obvious one (see e.g. Lehtonen et al., 2012). Thus, an asexual lineage invading the sexual population would be expected to quickly drive its sexual conspecifics extinct (Maynard Smith, 1978). None of these disadvantages apply to all sexual species because of the highly variable nature of their reproduction. But, under many circumstances, the disadvantages apply profoundly (Lehtonen et al., 2012). Thus, sexual reproduction remains an enigma that calls for explanation.

### Theories of sexual reproduction

Many main concepts and their countless variants were proposed to explain the paradox of sexual reproduction (reviewed e.g. in Williams, 1975; Maynard Smith, 1978; Bell, 1982, 1985; Kondrashov, 1993; Meirmans & Strand, 2010). The genetic advantages of sexual reproduction as the explanation for the maintenance of sexual reproduction are highlighted by theories like the Weismann’s idea of sex generating variability, later delimited as the concept of Vicar of Bray (Bell, 1982), Fisher-Muller’s accelerated evolution of sexual species (Muller, 1932; Fisher, 2003), breaking free of neighbouring deleterious mutations (Crow, 1970), reduction of the spread of genomic parasites (Sterrer, 2002), advantage of diploidy (Lewis & Wolpert, 1979), repair of DNA (Bernstein & Bernstein, 2013), restoration of epigenetic signals (Gorelick & Carpinone, 2009), eventually stochastic and deterministic variants of Muller’s ratchet hypothesis (Muller, 1964; Kondrashov, 1982).

Ecological theories of sexual reproduction, on the other hand, stress the assumption that sex provides some ecological advantage to sexual species. Certain trends can be clearly found in the geographic distribution of sexual reproduction, as was recently summarized by Hörandl (2006, 2009) or Vrijenhoek & Parker (2009). As most advantages of sexual reproduction postulated by genetic theories could be achieved by automixis (Gorelick & Carpinone, 2009; Neiman & Schwander, 2011), the correct answer to the “greatest paradox of evolutionary biology” probably lies in the group of ecological theories of sexual reproduction, or at least in some form of theoretical synthesis that includes them (West et al., 1999).

### Ecological theories of sexual reproduction and their predictions

Ecological theories such as Red Queen theory (Hamilton et al., 1990), evolutionary arm-races hypothesis (Dawkins & Krebs, 1979), and the fast-sexual-response hypothesis of Maynard Smith (1993) emphasize the sexually reproducing organisms’ advantage when interacting with other organisms, which are able to dynamically react in a co-evolutionary manner. According to these “biotic heterogeneity advantage” theories, sexual species should prosper in spatially and temporally biotically heterogeneous environments, i.e. environments with many biotic interactions from competitors, predators, and parasites – e.g. tropical rainforests, low-latitude coral reefs, ancient lakes, climax communities, or generally species-rich ecosystems In such environments, selective pressures affecting the offspring differ profoundly from those that affected the parents because of the coadaptation (or rather counter-adaptation) between interacting organisms. In the presence of such intensive biotic interactions, sexual species would be especially favoured because they maintain high genetic polymorphism and could quickly react to the countermoves of their evolutionary opponents by a simple change of allele frequencies in the population. The speed, not the depth, of adaptation is essential in these environments (Maynard Smith, 1993).

Another group of ecological theories of sexual reproduction sees the main advantage of sexual reproduction in the higher fitness sexual individuals or species have in abiotically heterogeneous environments. This group of “abiotic heterogeneity advantage” theories comprises, for example, the lottery and Sisyphean genotypes hypothesis (Williams, 1975), elbow room hypothesis (Maynard Smith, 1978), tangled bank hypothesis (Bell, 1982), hypothesis of fluctuating selection (Smith, 1980), and hypothesis of reduced response to fluctuating selection (Roughgarden, 1991). Such environments are abiotically very variable in space and/or time, i.e. diverse, unpredictable, and with unequally distributed resources – e.g. temporary or ephemeral habitats, exposed habitats, dynamically changing freshwater environments, coastal habitats or more generally biomes of high latitudes and/or altitudes. The offspring usually inhabit an environment different from that of their parents due to the dispersal in time and/or space. An abiotic environment does not co-evolutionarily react to the evolutionary moves of its inhabitants, allowing them to perfectly adapt to it under certain circumstances, e.g. under conditions of slow, long-term changes. Under these circumstances, the asexual species might have an advantage because, for example, they do not suffer from segregation and recombination loads (Crow, 1970). However, if the environment is very heterogeneous and unpredictable in time and/or space, the sexual species have the same advantage as in the biotically heterogeneous environment.

The heterogeneity of the environment, both biotic and abiotic, can be comprehended as the sum of heterogeneity in space (in the sense of variability) and time (in the sense of instability, especially when the change is unpredictable). The temporal and spatial aspects of heterogeneity, even though differing substantially at first sight, could act remarkably similarly in terms of favouring sexual species (Kondrashov, 1993; Otto & Lenormand, 2002; Neiman & Schwander, 2011). In principle, the most important factor is always whether the environment inhabited by the offspring differs in its character (i.e. selective pressures) from the environment inhabited by their parents. There are two conditions under which such a situation is fulfilled; the environment either changes dynamically in time, or is very patchy and the offspring often live in different microhabitats.

Both spatial and temporal heterogeneity could be the consequences of both biotic and abiotic factors (Li & Reynolds, 1995). Thus, according to the major source of the environmental heterogeneity, it is essentially possible to differentiate between “biotic” and “abiotic” ecological theories of sexual reproduction, despite the fact that the biotic and abiotic parts of the environmental heterogeneity are usually interconnected and that they influence and complement each other in their effects on the advantage of sexual or asexual reproduction (Glesener & Tilman, 1978). Abiotically homogeneous environments can (in the case of plentiful resources and mild conditions), but do not have to (e.g. in the case of extreme environments) cause an increase of the biotic aspect of environmental heterogeneity (e.g. by increasing the complexity of the ecosystem) (Tokeshi, 1999). Abiotic heterogeneity could act both ways too. It might cause an increase of biotic heterogeneity of the environment, or at least of the species diversity. On the other hand, abiotically heterogeneous environments with conditions widely fluctuating to extremes (e.g. the desiccating ponds) tend to be biotically homogeneous (Tokeshi, 1999).

The “biotic” and “abiotic” theories of sexual reproduction mentioned above have different predictions regarding the character of the environment that will be advantageous for sexual and asexual species. However, their predictions are not absolutely disparate—one could easily devise examples of environments suitable for asexual species according to both groups of theories, e.g. stable extreme environments. Similarly, the individual theories of sexual reproduction are far from being disparate; they are usually interconnected in their basic principles, they intermingle and complement each other (Meirmans & Strand, 2010). Many, if not all “abiotic” theories can be reinterpreted “biotically”. It is therefore possible that the differentiation of “biotic” and “abiotic” ecological theories is important in theory, but not important in the real world, and that sexual organisms do have an advantage in environments that are both biotically and abiotically relatively heterogeneous (i.e. overall heterogeneous environments).

It was suggested by Williams (1975, pp. 145–146, 149–154, 169) and explicitly discussed by Flegr (2013) that the lower ability of sexual species to fully adapt to transient environmental changes brings them, paradoxically, a major advantage in randomly fluctuating environments, i.e. in environments expressing large (biotic or abiotic) heterogeneity in time. Evolutionary “plastic” (*sensu* Flegr) asexual species, sooner or later, adapt to transiently changed environmental conditions and become extinct when the conditions return to normal. Sexual species are not able to fully adapt to temporal environmental changes; they always retain some genetic polymorphism that helps them escape extinction when the conditions quickly return to normal. This hypothesis was supported by the results of certain experimental studies, e.g. long-term patterns of fitness and genetic variability (Renaut et al., 2006) or dynamics of adaptation (Colegrave et al., 2002; Kaltz & Bell, 2002) in sexually and asexually reproducing *Chlamydomonas*. According to this (meta-)hypothesis, asexuals would prevail in stable or predictively slowly changing, possibly extreme, environments of low temporal heterogeneity (Flegr, 2013). Given the similarities between the effect of temporal and spatial heterogeneity mentioned above, this notion can be readily extended to encompass both temporal and spatial heterogeneity. Similar remarks were made, e.g. by Williams (1975, p. 153) and Roughgarden (1991), while a combination of several aspects of heterogeneity was implicitly also proposed as the explanation of the presence of sexual reproduction by Glesener & Tilman (1978) and some interpreters of the Red Queen theory (e.g. Butlin et al., 1999) or the tangled bank hypothesis (e.g. Bell, 1982).

### Comparing the ecology of sexual and asexual groups

Most of the organisms that live on Earth, Archaea and Bacteria, are primarily asexual, while most of the known species, eukaryotes, are primarily sexual (Speijer et al., 2015). However, some eukaryotic lineages switched to secondary asexual reproduction (de Meeus, Prugnolle & Agnew, 2007; Van Dijk, 2009; Speijer et al., 2015). It is therefore possible to compare the environmental biotic and abiotic heterogeneity of such secondary asexual clades with that of their sexual relatives.

Various studies aimed at testing and discriminating between individual ecological theories of sexual reproduction on the basis of their predictions about the environmental correlates of sexual and asexual lineages show largely inconclusive results. Often their aim was to test particular theoretical concepts: lottery hypothesis and Sisyphean genotypes hypothesis (Williams, 1975; Hörandl, 2009), elbow room hypothesis (Garcia & Toro, 1992; Koella, 1993), Red Queen theory (Burt & Bell, 1987; Neiman & Koskella, 2009), fast-sexual-response hypothesis (Becerra et al., 1999), hypothesis of optimal responsibility to fluctuating selection (Griffiths & Butlin, 1995; Schön & Martens, 2004), hypothesis of prevention of loss of genetic variability under fluctuating selection (Maynard Smith, 1993; Hörandl, 2009; Vrijenhoek & Parker, 2009) or tangled bank hypothesis (Vrijenhoek, 1984; Burt & Bell, 1987; Griffiths & Butlin, 1995; Domes et al., 2007; Maraun et al., 2012); or at least they were later interpreted as such. The most extensive comparison not focused on testing one particular theoretical concept was performed by Bell (1982) on multicellular animals (Metazoa). In his comprehensive study, he concluded that the results mostly supported the tangled bank hypothesis. However, his study included both old and young asexual taxa and might not fully appreciate the influence of different life strategies on the character of the environment the species subjectively experience— see the Discussion. Experiments aimed at discriminating the selective pressures of biotically (see e.g. Fischer & Schmid-Hempel, 2005) or abiotically (see e.g. Becks & Agrawal, 2010) heterogeneous and homogeneous environments were also performed, mostly pointing to the conclusion that heterogeneous environments select higher rates of recombination or sexual reproduction. However, particular mechanisms that favour higher levels of sex are hard to determine in these cases that are, moreover, often based on facultatively sexual organisms.

### Short-term and long-term asexual groups

The main problem of the comparative studies mentioned above may be the inclusion of both old and young asexual taxa. Most secondary asexual groups probably are not evolutionarily viable in the long term, as could be deduced from the distribution of asexual lineages on the “tree of life.” With the exception of several ancient asexuals (AAs), they form only the terminal twigs—species and genera (Butlin, 2002). This pattern could be the consequence of the disadvantage that secondary asexual species have in species selection (Nunney, 1989). In the large majority of cases, young asexuals do not speciate, and when the conditions suitable for their opportunistic transition to asexuality change, they become extinct. Another suggested reason for this macroevolutionary failure of asexual species is their failure in the interspecific sorting in fluctuating environments—asexual species are able to adapt to temporarily changed conditions more easily, but they fail to readapt quickly enough because of the lack of genetic polymorphism, and therefore become extinct when the conditions return back to normal (Williams, 1975; Flegr, 2013). At least some young asexual lineages could, in fact, consist of short-lived clones continuously cleaved from maternal sexual population (Janko et al., 2008; Vrijenhoek & Parker, 2009). Alternatively, they could be sustained by an occasional hybridisation with related sexual lineages (Turgeon & Hebert, 1994; van Raay & Crease, 1995; Butlin et al., 1998b) or an infrequent transfer of genetic material from “host species” in hybridogenetic and gynogenetic lineages (Mantovani et al., 2001; Bogart et al., 2007). In sum, young asexuals do not have to exhibit the properties that would allow them to survive in the long term and, contrary to the mainstream view, they could in fact bring a significant noise into the studies of long-term maintenance of sexual (and secondary asexual) reproduction. In other words, the existence of most asexual lineages might be a consequence of certain ecological opportunism and the reasons for their survival might be, in contrast to the AA lineages, different from case to case. We speculate that this may be the main reason for the inconclusiveness of most comparative studies mentioned above—they focused only on short-term asexual clones or did not take into account the age of different clones.

The phenomenon of asexual “terminal twigs contra ancient asexuals” is still somewhat controversial and its real existence is being discussed (see e.g. Schön et al., 1996; Neiman et al., 2009; Schwander & Crespi, 2009; Janko et al., 2011). It was even suggested that the cumulative frequency distribution of ages of asexual lineages is “fairly regular” and nearly linear on the logarithmically transformed scale, i.e. it shows no obvious break between the young and old asexuals (Neiman et al., 2009). As a consequence, AAs need not be substantially different from other asexuals and their delimitation could be more or less arbitrary. On the other hand, Janko et al. (2011) strongly questions the methodology of Neiman et al. (2009), especially the choice of linear regression on logarithmic scale and its interpretation. They argue that the data distribution presented by Neiman et al. (2009) is unlikely to provide any strong conclusions regarding the universality of processes affecting clonal life spans—in fact, it fits even better to other distributions, including a multidiffonential distribution, which is characteristic of distributions generated by a mix of different underlying processes. Moreover, using their analysis of clonal genetic variability, Janko et al. (2011) identified a growing selective pressure against asexual lineages based on their age, corroborating the hypothesis of the long-term disadvantage of asexuality. Regardless of these discussions, it is obvious that out of all the secondary asexual clades only the AAs have been able to survive or even diversify in an asexual state for millions of years (Judson & Normark, 1996; Normark et al., 2003; Neiman et al., 2009; Schurko et al., 2009; Schwander & Crespi, 2009). This is the main reason that our study is based exclusively on AAs as they already proved to be evolutionarily viable in the long term. Contrastingly, young opportunistic clonal lineages still face a serious extinction risk during the first pronounced change of conditions.

However, it is worth mentioning that the focus on AAs puts forward another serious difficulty: these clades were separated from their sister sexual lineages a long time ago (at least 1 million years ago, see Materials and Methods) and both sexual and asexual lineages thus underwent considerable time periods of independent evolution. Therefore, both lineages independently acquired numerous adaptations that distinguished them but need not be related to the mode of their reproduction. Singular case studies comparing AAs and their sexual sister lineage thus are not expected to have a strong predictive value in the long-term maintenance of asexual reproduction. On the other hand, an extensive comparative study would enable the comparison of several such pairs of AA and sexual control and reveal possible common adaptations of AAs related to their long-term survival in an asexual state.

### Aims of the study

The main aim of the current study is to test whether AAs more often inhabit 1) generally less heterogeneous environments, 2) less biotically heterogeneous environments, or 3) less abiotically heterogeneous environments. To this end, we used paired exact tests to compare the ecological demands of sexual species and AA species within all six unrelated clades of eukaryotic organisms in which the presence of differences in associated environmental heterogeneity can be recognised on the basis of available published data. Namely, we tested the hypothesis that, within such pairs, the asexual species inhabit more homogenous environment more often than the sexual species. In the exploratory part of the study, we searched for particular environmental properties and organismal adaptations that are common among the AA members of the pairs.

## Materials and Methods

### Identification of ancient asexuals and their sexual controls

#### Ancient asexual groups

The definition of the “ancient asexual group” is rather vague. Some researchers consider a lineage to be AA if it reproduces obligately asexually for at least 50 000 generations or 0.5 million years (Law & Crespi, 2002); some prefer one million generations (Schwander et al., 2011), yet others just speak about “millions of years” (Judson & Normark, 1996; Normark et al., 2003). It was even suggested that AAs are not substantially different from other asexuals and their delimitation is more or less arbitrary (Neiman et al., 2009). It is not the aim of this study to argue for the substantial difference of AAs from other asexuals or against it. We focus only on groups that were proven to survive in an asexual state for a considerable amount of time. Thus, regardless of the discussion on the fundamental distinction of young and old asexual taxa, which was briefly presented in the Introduction, in the current study we defined AAs conservatively as those secondary asexual eukaryotic lineages that reproduce obligately asexually with a great deal of certainty for at least one million years.

The condition of obligate asexuality is at least equally as important as the age of the studied asexual groups in the context of this study, as many seemingly asexual lineages might experience rare sexual events or other forms of genetic exchange. These options constitute a greater problem in short-term asexuals than in AAs, which are spatially and temporally isolated from their sexual sister lineages. Nevertheless, their obligate asexuality was discussed, studied, and verified thoroughly (see Table S1).

At the beginning, we identified well supported AA groups with the help of literary sources. We started with published secondary literature such as Judson & Normark (1996), Normark et al. (2003), Neiman et al. (2009), Schurko et al. (2009), Schwander & Crespi (2009) and Speijer et al. (2015), investigated cited primary literature and other novel primary literal sources concerning putative AA groups. We also investigated other possible AAs proposed in the primary literature and some lineages traditionally believed to be long term asexual. The evidence for confirmation or rejection of putative AAs included organismal, life-history, palaeontological, biogeographical, molecular, individual genetic, and population genetic data and also other indices of ancient asexuality proposed in the AA literature listed above. The list of supported and contested AA candidates, as well as reasons for our decision, are summarized in Table S1. The confirmed AA groups were eight: bdelloid rotifers (Bdelloidea), darwinulid ostracods (Darwinulidae), several lineages of oribatid mites (Oribatidae), several lineages of mites from the suborder Endeostigmata and order Trombidiformes, shoestring fern *Vittaria appalachiana* (Farrar & Mickel), three species of stick insects from the genus *Timema*, and several lineages of bivalve genus *Lasaea*. Only these groups were included in our comparative study.

#### Sexual controls

In the next step, we identified ecologically comparable sexual sister lineages for the eight AA groups by using literary sources. In those individual cases in which the phylogenetic relations between the sexual and asexual lineages were not entirely clear, we used the closest possible comparable clades—clades proven to be closely related and comparable in terms of their ecology (see Table S2). Three of the AA groups were monophyletic (Bdelloidea, Darwinulidae, and *Vittaria*). The remaining AA groups were polyphyletic, i.e. they included several related monophyletic asexual sub-lineages with interstitial sexual lineages. In each case of a polyphyletic AA taxon, these closely related AA lineages were taken together and treated as single unit in the analysis. In the opposite case, i.e. if the monophyletic lineages of these polyphyletic taxa were incorporated into the analysis as single units, the risk would arise that the observations would not be independent and the results of statistical analysis could be biased because of the effect of pseudoreplications. Thus, in these cases, we compared every individual AA lineage with its sister sexual lineage in the monophyletic sub-taxa of the polyphyletic AA groups and based our conclusions on the prevailing trend (i.e. over 50 % of the cases; however, all actual trends were much more convincing, see Table S3) in the polyphyletic groups. Unfortunately, with the exception of *Timema*, the internal phylogenetic relationships of the studied polyphyletic AA groups were more or less unclear. Where possible, we proceeded using the most probable relationships (Bdelloidea, Darwinulidae, Oribatidae, Nematalycidae and Proteonematalycidae, Grandjeanicidae and Oehserchestidae, see Table S2). In the cases with several equally probable alternative phylogenetic relationships of AA and sexual lineages (both in monophyletic /*Vittaria/*, and polyphyletic /*Alicorhagia* and *Stigmalychus*, Pomerantziidae, *Vittaria, Lasaea/* AA taxa), we compared AA lineages with alternative sexual controls to determine the consistency of the trend in the association of AA lineages or sexual controls with biotically and/or abiotically more heterogeneous environments (all trends were consistent over all alternative sexual controls, see Table S3).

It could be argued that it is legitimate to consider particular asexual monophyletic lineages in the polyphyletic AA group as independent observations when we inquire into the reasons for maintaining sexual reproduction, not into the reasons for the transition to sexuality (Thornhill & Fincher, 2013). However, we chose the more conservative approach because, as noted above, the phylogenetic relationships of the individual AA and sexual lineages were often not clear enough to form sisterly pairs of sexual and asexual lineages in the polyphyletic AA groups with necessary certainty.

#### Determination of environmental heterogeneity

After the identification of the AA groups and their sexual controls, we focused on the comparison of the character of their environments. Using relevant literary resources, we collected and analysed data on the (biotically or abiotically more heterogeneous or homogeneous) character of environments inhabited by the studied groups (see Table S3). Biotic and abiotic environmental heterogeneity clearly have a non-trivial relationship to each other (see the Discussion), but it is essentially possible to distinguish them. Environmental heterogeneity, both biotic and abiotic, is an emergent property stemming from different factors and different adaptations in various AAs and thus could not be quantified and rated on a single universal scale. On the other hand, there is no obstacle of comparing AAs and their sexual controls based on this general property of their environment.

#### Biotic heterogeneity

We define biotically highly heterogeneous environments as those in which selective pressures affecting the offspring differ profoundly from those that previously affected their parents because of the coadaptation (or rather counter-adaptation) of interacting organisms. Thus, the most biotically heterogeneous environments are the habitats with a high degree of competition, predation, and parasitism. Biotic heterogeneity has both temporal (the coadaptation of interacting organisms) and spatial (e.g. the migration of the offspring to the areas with new competitors, predators, and parasites) dimensions. Changes in the biotic heterogeneity are essentially unpredictable, with the exception of some ecological cycles (e. g. host-predator or host-parasite cycles). In the latter case, environments with unpredictable changes were considered more biotically heterogeneous.

For example, the environment of organisms that live in tight association with other organisms is biotically very heterogeneous. This applies especially to predators and parasites that are forced to respond to the evolutionary counter-moves of their prey and hosts (Dawkins & Krebs, 1979). The more specific relationship with prey and hosts they have, the stronger the selective pressures of counter-adapting prey and hosts affect them (Dawkins & Krebs, 1979). Therefore, it is expected that organisms that use non-specific predatory strategies, e.g., filtering (especially if they filter both living organisms and dead organic matter), are under relatively weak selective pressure from their prey. Their environment is consequently biotically relatively homogenenous in this regard. On the other hand, the environment of organisms that are themselves under strong selective pressure of predators and parasites is highly biotically heterogeneous (Dawkins & Krebs, 1979). The environemnt of organisms that are not under strong selective pressures of predators and parasites for various reasons is biotically more homognenous in this regard (e.g. the environment of Darwinulidae, see Schön et al., 2009b or Bruvo et al., 2011). Another important component of environmental biotic heterogeneity is competition. The environments with complex ecosystems that are characterized by high a degree of competition, predation and parasitism among their inhabitants (e.g. ancient lakes, see Martens, 1998; Martens & Schön, 2000; Schön & Martens, 2004) are highly biotically heterogeneous for them. On the other hand, the environments with a low degree of competition, predation and parasitism (especially extreme habitats, e.g. the environments of extremely high temperatures, see Tobler, 2007, or, for photosynthetic organisms, poorly lit environments, see Farrar, 1978, 1998), are biotically very homogeneous for their inhabitants. The vast majority of environments on Earth are thus somewhat biotically heterogeneous. In spite of that, we can find important exceptions. This factor of environmental heterogeneity is considerably weakened especially in extreme, ephemeral and marginal habitats that cannot sustain complex ecosystems because of their extreme conditions, rapid unpredictable changes, low carrying capacity and/or insufficient energy sources (see e.g. Bell, 1982).

An important factor that affects the biotic heterogeneity of environment the organisms experience is the way of life practiced by aquatic organisms. On the average, lesser biotic heterogeneity is experienced by benthic (or sedentary) organisms in comparison with planktonic aquatic organisms. The reason is that the latter are subject to fast and effective transmission of parasites and pathogens, especially viruses (see e.g. Suttle et al., 1990; Bratbak et al., 1993; Fuhrman, 1999; Wommack & Colwell, 2000; Suttle, 2005, 2007), because of the character of their environment—mixing of water masses lead to frequent encounters of various individuals (Emiliani, 1993a, b). Crucial difference of the resulting risks for benthic and planktonic organisms was pointed out by Emiliani (1982; 1993a). Emiliani (1993a) documented that benthic representatives of Foraminifera have lower risk of extinction in comparison with planktonic ones. The average length of existence of their benthic species was 20 million years, whereas planktonic species lasted only about 7 million years. The most probable explanation of this pattern is higher susceptibility of planktonic organisms to extinction caused by lethal parasitic, especially viral, infections. This finding gave rise to the viral theory of background extinctions (Emiliani 1993a, b). Evidence for the lower risks arising for benthic organisms from parasites and pathogens was also supported by other ecological studies. For example, Filippini et al. (2006) observed lower prevalence of individuals infected by viruses and consequent mortality among benthic bacteria (~0,03 %) in comparison with bacteria from water column (~6 %). Moreover, this pattern held despite much larger abundance of viruses in the sediment of studied temperate lake. Fisher et al. (2003) and Bettarel et al. (2006) came to very similar conclusions on the basis of studies of temperate oxbow lake and freshwater habitats in tropical Africa, respectively (but see Danovaro et al., 2008 for somewhat contrasting results from marine benthic sediment). Putting aside the limitations of parasite and pathogen transmission among benthic organisms, the protection of benthic organisms against viruses might be further enhanced by the adsorption of viruses into organic and inorganic particles of sediment and the aggregation of benthic organisms (Filippini et al. 2006; Fisher et al., 2003). Whereas the extinction of whole species due to viral infection (eventually infection by another parasite or pathogen, e.g. fungus) is possible only under very limited conditions (see e.g. Buckwold, 1994, or de Castro & Bolker, 2005), local extinctions caused by pathogen or parasitic infections are probably quite common (Emiliani, 1982, 1993a; de Castro & Bolker 2005). The planktonic way of life thus probably considerably increases the selective pressures of parasites and pathogens and consequently the biotic heterogeneity of the environment.

The argument for lower biotic heterogeneity of benthic (or sedentary) organisms is possible to extend also to organisms that inhabit soil. From the viewpoint of the viral theory of background extinctions (Emiliani, 1993a, b), soil represents an environment that considerably impedes the spread of pathogens and parasites. Interactions with parasites and pathogens are very limited both in intensity and frequency due to the tortuous, i.e., multidimensional, character of soil matrix—it is best described as semi-discontinuous network of pores filled with air and/or water, or water films surrounding solid particles (Lavelle & Spain, 2003; for more information about the character of soil environment see also Wallwork, 1970; Coleman et al., 2004, or Paul, 2007). These features of soil matrix limit both the passive spread of parasites and pathogens and the frequency of encounters among their transmitters and organisms in general. The tortuous character of soil environments was stressed as a factor that limits the passive spread of viruses in benthic sediment by Fisher et al. (2003), whereas Murphy & Tate (1996) emphasized its limitations on the spread of bacteria. These observations are in agreement with Drake et al. (1998), who observed a negative correlation between the concentration of viral particles and sediment grain size. The sieving effect of the soil for organisms of various sizes is also commented by Paul (2007). Moreover, the tortuous character of soil impedes also active dispersion of organisms, e.g. when searching for prey (Elliott et al., 1980). Only a few larger organisms are able to effectively move larger distances within soil or even create their own habitats; movements of most organisms are locally constrained (Lavelle & Spain, 2003). Direct encounters between organisms, even organisms of the same species, are thus relatively infrequent. This leads, together with the limited spread of pheromones (Karasawa & Hijii, 2008), for example, to frequent transitions to indirect fertilisation with the help of deposited spermatophores (see e.g. Wallwork, 1970, or Lavelle & Spain, 2003). The pattern of spatial autocorrelation of genetic lineages in soil communities, for example in rotifers (Robeson et al., 2011), further supports the limited dispersal abilities of soil organisms. The genetic diversity of various Bdelloidea lineages in soil is correlated only on small spatial scales (up to 54–133 m). Operational taxonomic units identified by Robeson et al. (2011) almost did not overlap above this distance. Habitats that distanced only tens to hundreds of meters were thus inhabited overwhelmingly by separate genetic lineages. Moreover, rotifer communities differed to a certain degree even in the smallest investigated distance of 16 cm (Robeson et al., 2011)^1^.

Taken together, the tortuous character of the soil affects all soil organisms at various scales not only in terms of the reduced spread of parasites and pathogens, but also lower frequencies of encounters with predators and competitors. This leads to an overall reduction of biotic pressures in soil, which is further supported by the striking evolutionary stasis of many lineages of soil inhabiting organisms (Pilato, 1979). Moreover, species richness and population sizes, including parasites, predators and competitors, markedly decreases with the depth of soil horizon (Lavelle & Spain, 2003; Paul, 2007). Deep soil horizons are therefore even more abiotically homogeneous. The specific character of soil environment does not imply its general spatial homogeneity. On the contrary, soil is often spatially heterogeneous, especially on a larger scale (see e.g. Lavelle & Spain, 2003; Coleman, Crossley & Hendrix, 2004, or Paul, 2007). It is the tortuous and multidimensional character of soil matrix that reduces biotic pressures affecting its inhabitants and makes this environment biotically very homogeneous.

The biotic heterogeneity of the environment the organisms experience might be reduced by the presence of durable resting stages. Organisms may get rid of parasites and pathogens, survive unfavourable environmental conditions, or colonize new habitats with naïve parasites, predators, pathogens and competitors in these stages (as do, for example, Bdelloidea—see Wilson, 2011). The geographical trend of decreasing biotic heterogeneity with increasing latitude might be expected on a global scale. Species diversity and ecosystem complexity decrease with distance from equator (Tokeshi, 1999). These events are coupled with a decreasing intensity of parasitization, abundance, prevalence and a relative diversity of parasites (Rohde, 1986; Rohde & Heap, 1998). An analogous trend of decreasing biotic heterogeneity with increasing depth might be expected in deeper parts of the water column for the same reasons (see e.g. Etter et al., 2005).

#### Abiotic heterogeneity

Abiotically highly heterogeneous environments are defined as those that are highly variable regarding changes of abiotic factors. They are diverse, unstable, and have unequally distributed resources. Again, the abiotic heterogeneity of the environment has both spatial (in the sense of variability) and temporal (in the sense of instability) dimensions. The offspring thus usually inhabit an environment different from that of their parents due to their dispersal in time and/or space. Changes in the abiotic environment could be predictable (e.g. cyclical) or unpredictable, and their intensity and frequency vary on different timescales. We are interested in ecological timescales in this study so we consider short-term unpredictably changing environments as the most abiotically heterogeneous.

Temporally and spatially highly changeable ephemeral and marginal habitats are especially abiotically heterogeneous environments (see e.g. Pejler, 1995). However, most of the surface terrestrial habitats are considerably abiotically heterogeneous. On the contrary, sheltered habitats such as caves, ground water reservoirs or soil environment are greatly abiotically homogenenous. Such environments protect their inhabitants from solar radiation and buffer short-term fluctuations in outer environment (e.g. changes of temperature and humidity), protecting their inhabitants from the direct impacts of such changes (Wallwork, 1970; Farrar, 1978, 1990, 1998; Krivolutsky & Druk, 1986; Lavelle & Spain, 2003; Devetter & Scholl, 2014). Most changes in soil matrix are much slower in comparison with surface habitats (Lavelle & Spain, 2003). The abiotic homogeneity of soil environment further increases with the depth of the soil horizon. For example, there is a specific depth of soil horizon in each geographical region under which the temperature is perienally stable, depending on its latitude, altitude and other climatic factors (Wallwork, 1970; Lavelle & Spain, 2003; Coleman et al., 2004; Paul, 2007). The buffering effect of soil on moisture fluctuations also increases with depth (see e.g. Quesada et al., 2004). Moreover, soils of certain biomes (especially forest soils) are temporally abiotically more homogeneous than soils of other biomes (see e.g. Siepel, 1994, 1996).

Regarding aquatic environments, freshwater habitats and coastal areas are the most abiotically heterogeneous (Sheldon, 1996). The decrease of abiotic heterogeneity with increasing depth is also expected—water masses buffer surface environmental changes in a similar way to soil (see e.g. Etter et al., 2005). Certain extreme environments that are temporally stable (e.g. hot springs or subsurface cavities) are also very abiotically homogeneous (Bell, 1982), but this does not apply to all environments referred to as extreme.

In a similar way to the biotic heterogeneity of the environment, also the abiotic one might be reduced in the populations of organisms producing durable resting stages. Such an adaptation enables them to survive unfavourable fluctuations of an abiotic environment and promotes colonization of new habitats (see e.g. Wilson, 2011). On the other hand, mobility probably does not strongly affect the abiotic environmental heterogeneity that the organisms experience. Mobile organisms might hypothetically experience more different abiotic conditions in their life, but they can also easily stick with those most suitable for them. The geographical trend of increasing abiotic heterogeneity with increasing latitude and altitude might be expected to occur on global scale (Hörandl, 2006, 2009; Vrijenhoek & Parker, 2009). However, it is noteworthy that such a trend might be countered by an opposite trend of the decreasing biotic heterogeneity mentioned above in its effects on sexual and asexual species (and *vice versa*). All of the expectations mentioned above need not apply absolutely, but may serve as useful leads in judging the environmental heterogeneity of various organisms if their peculiarities are taken into account.

In two AA groups (*Lasaea, Timema*) we were unable to identify any consistent differences in the heterogeneity of the environments inhabited by their sexual and asexual lineages. The probability that any two groups experience absolutely identical habitat heterogeneity is negligible and the absence of difference must be just a result of lack of empirical data. *Lasaea* and *Timema* were thus not included in the statistical analysis because the binomial test can analyse only binary variables (see e.g. McDonald, 2014) and our current knowledge on these groups’ ecology does not allow us to decide whether asexual or sexual species live in more heterogeneous environments. The same applies for the abiotic heterogeneity of the environment of Darwinulidae. Thus, the analysis was based on only six pairs of AAs and their sexual controls, out of which six differed in the abiotic heterogeneity of their environment and five in the biotic heterogeneity.

#### Statistics

Collected data were analysed using the R v. 3.1.2 software environment (R_Core_Team, 2014). We chose to evaluate data by an exact test, specifically a one-tailed binomial test, due to the low number of compared sexual-asexual pairs (six pairs in total, six for biotic heterogeneity and five for abiotic heterogeneity), the character of the data (pair, binary), and the type of the tested hypothesis. By this means, we tested three hypotheses, namely that AA groups inhabit predominantly (1) biotically or abiotically, (2) biotically, and (3) abiotically more homogeneous environments than their sister or closely related ecologically comparable sexual groups.

## Results

### Ancient asexuals

Reviewing relevant literary resources, including several recent studies, we concluded that at least eight of the putative AA groups do fulfil our strict criteria of ancient asexuality: bdelloid rotifers (Bdelloidea), darwinulid ostracods (Darwinulidae), some mite lineages of the Oribatidae, Endeostigmata, and Trombidiformes, shoestring fern *Vittaria appalachiana*, some lineages of stick insects from the genus *Timema*, and some lineages of the bivalve genus *Lasaea*; see Supplemental table S1. Their sister or closely related ecologically comparable sexual groups were identified consequently with the help of relevant literature; see Table S2.

### Comparative analysis of the habitat homogeneity of ancient asexuals and their sexual controls

The comparison of the character of environments inhabited by the AAs and their sexual controls showed that AAs inhabit biotically or abiotically (6 of 6, p = 0.016), biotically (6 of 6, p = 0.016), and abiotically (5 of 5, p = 0.031) more homogeneous environments. All these results are statistically significant. Details of the results are summarized in the Supplemental review of AA ecology and Table S3.

### Exploratory part of the study

The exploratory part of this study is based on data gathered in the course of the main analytic study. We identified several properties and adaptations that are common to a considerable number of studied AAs. The most notable are durable resting stages, life in benthos and soil, and life in the absence of intense biotic interactions. These findings are discussed below.

## Discussion

In contrast with other comparative studies in the field, the presented one is (1) based exclusively on the AA taxa. Moreover, (2) biotic and abiotic environmental heterogeneity has been strictly distinguished. We conclude that all 6 of the 6 analysed AA groups inhabit biotically more homogeneous environments and all 5 of the 5 ones inhabit abiotically more homogeneous environments when compared with their sexual controls. Some of the studied AA groups are found in environments of a predominantly low biotic homogeneity (mainly Darwinulidae) or a predominantly low abiotic homogeneity (mainly Oribatidae), but most of them are associated with environments substantially more homogeneous both biotically and abiotically (Bdelloidea, non-Oribatidae mites, *Vittaria*). No AA group lives in an environment abiotically or biotically more heterogeneous than its sexual control. In the cases excluded from the analysis (abiotic heterogeneity in Darwinulidae and both biotic and abiotic heterogeneity in *Timema* and *Lasaea*), it was not possible to distinguish whether the heterogeneity is lower in the AA group or in the sexual control. More specifically, enough information could not be found to decide either way.

The associations with biotically and abiotically more homogeneous environments overlap almost perfectly. Thus, the results of the comparative analysis clearly indicate that either the AA groups tend to be associated with overall (both biotically and abiotically) homogeneous environments, or that these two types of heterogeneity are so strongly correlated that it is impossible to decide in favour of theories of sexual reproduction that stress the key role of biotic or abiotic heterogeneity. In general, our results obtained on AAs support, but of course do not prove, the hypotheses that consider both biotic and abiotic heterogeneities acting as one factor in their effect on organisms (Williams, 1975, pp. 145–146, 149–154, 169; Roughgarden, 1991; Flegr, 2010, 2013).

Despite the widespread apprehension that the long independent evolution of AAs and their sexual controls would hamper any ecological comparative analysis of the type presented here (leading to the preference of young asexual lineages, see Introduction), we found that both groups usually inhabit quite similar and considerably homogeneous environments. Contrary to the initial expectation, this can, in fact, complicate analyses aimed at evaluating the differences of the environments of AAs and their sexual controls in the opposite way (as was the case of *Timema* and *Lasaea*, see Table S3). On the other hand, their common ancestor’s association with the homogeneous environments could have been a preadaptation to the successful and long-term transfer to asexual reproduction in the AAs. This tendency is obvious especially in Darwinuloidea-Cypridoidea, but it can also be seen in Bdelloidea-Monogononta, Oribatidae, and Endeostigmata (see Table S3).

It is interesting in this regard that a lot of contested AAs (see Table S1) also inhabit considerably homogeneous environments—e.g. arbuscular mycorrhizal fungi of the order Glomales (Croll & Sanders, 2009), tardigrades (Pilato, 1979; Mobjerg et al., 2011), nematode genus *Meloidogyne* (Castagnonesereno et al., 1993), ostracods *Heterocypris incongruens* (Ramdohr) and *Eucypris virens* (Jurine) (Butlin et al., 1998b; Martens, 1998), bristle fern *Trichomanes intricatum* (Farrar) (Farrar, 1992), basidiomycete fungal families Lepiotaceae and Tricholomataceae (Currie et al., 1999a, b), ambrosia fungi Ophiostomatales (Farrell et al., 2001), or brine shrimp “*Artemiaparthenogenetica*” (Bowen & Sterling) (Vanhaecke et al., 1984)—and their adaptations are similar to those of the AAs included in this study (see below).

### What environmental properties and organismal adaptations are associated with AA taxa?

The exploratory part of this study is based on data gathered in the course of the analytic part of the study. Besides the tendency to inhabit biotically and abiotically homogeneous environments, we discovered several properties and adaptations that are common to a considerable number of studied AAs, do occur in AAs strikingly more often and could be the particular adaptations enabling their long-term survival in the environments mentioned above. Universally distributed adaptations potentially connected to the mode of reproduction were not expected to be found in our sample because of markedly different ecological strategies of the studied AAs. In spite of that, we found characteristics that have broad distribution among studied taxa:

### Alternative exchange of genetic information

Alternative ways of exchange of genetic information could theoretically substitute sexual reproduction and thus were repeatedly proposed as the key adaptation to asexuality (Butlin et al., 1998a; Gladyshev et al. 2008; Boschetti et al. 2011; Debortoli et al., 2016; Schwander, 2016). However, we identified this factor only once in the AAs included in our study (i.e. in 1/8 cases), namely in Bdelloidea that experience intensive horizontal gene transfer (Gladyshev et al. 2008; Boschetti et al. 2011; Debortoli et al., 2016). Another mechanism of genetic exchange, parasexuality (*sensu* Pontecorvo 1954), was proposed in some contested ancient asexuals— Glomales (Croll & Sanders, 2009), Tricholomataceae and Lepiotaceae (Mikheyev et al., 2006) and certain protists (Birky, 2009). However, considering only the well supported AAs, these mechanisms have limited distribution.

### Durable resting stages and subjectively homogeneous environment

There is one particular, mostly unnoticed adaptation that could be essentially important for the understanding of the links between the heterogeneity of environment and the mode of reproduction. It is based on the insight that the character of the environment is “read” much differently by its inhabitants than by the human observer. In case that a particular organism reacts to the adverse change of environmental conditions by entrenching itself in the resting or durable persistent stages (e.g. anabiosis) then, as a result, it *de facto* does not experience the unfavourable conditions at all. Its objectively heterogeneous environment becomes subjectively much more homogeneous.

This subjectivity addresses especially abiotic factors of environment, e.g. desiccation, which is survived in the anabiotic stages by Bdelloidea (Pilato, 1979; Ricci, 2001), or freeze and desiccation, which is survived in a state of torpor by Darwinulidae and some of their sexual relatives (Carbonel et al., 1988). Similar durable stages could also be found in some contested AAs, namely “*Artemiaparthenogenetica*”(Vanhaecke et al., 1984) and tardigrades (Mobjerg et al., 2011). Moreover, AA *Lasaea* is able to become mostly inactive and rests during the adverse conditions for some time as well (Morton et al., 1957). On the other hand, at least in Bdelloidea, the anhydrobiosis serves as the escape from biotic stresses too—especially parasites, both directly (the individual gets rid of parasites during desiccation) and indirectly (by enabling the escape from parasites in space and time) (Wilson, 2011). The distribution of durable resting stages among well supported AAs looks rather scarce (3/8 cases). However, these 2–3 groups comprise all studied AAs associated with significantly objectively abiotically heterogeneous habitats.

Despite the undeniable importance of this phenomenon and the fact that it was repeatedly highlighted, both historically (von Uexküll, 1982) and recently (Pilato, 1979), there is no general awareness of the idea of difference between the subjective environment experienced by the organism and the objective environment perceived by the human observer. This might be another reason why most researchers did not come to unambiguous conclusions in their comparative analyses of the ecology of sexual and asexual organisms. For example, a lot of “extreme” environments are not necessarily abiotically very homogeneous, whereas some environments that appear very abiotically heterogeneous on first sight (periodical ponds, dendrotelms etc.) could be very subjectively homogeneous for local inhabitants (e.g. anhydrobiotic Bdelloidea). After all, the “extremeness” of the environment itself depends on the adaptation of the observer, including the presence or absence of the durable stages, and thus is highly subjective.

For a species, the ability to survive adverse conditions entrenched in durable persistent stages could have crucial evolutionary consequences. It can be at least partially responsible for the phenomenon of evolutionary stasis, great morphological uniformity, and conservatism of numerous AA groups (see below) and indirectly responsible for the ability of successful and long-term transition to asexuality. Such organisms are shielded against the selective pressures of the environment. Their evolution is slow, but they are perfectly adapted to the conditions of their habitat (Pilato, 1979).

### Sedentary life and life in benthos

At least three well-supported AA groups (Bdelloidea, Darwinulidae, and *Lasaea*) are exclusively benthic or sessile in contrast to their sexual relatives (Dole-Olivier et al., 2000; Ricci & Balsamo, 2000; Ó Foighil, 1989). Some species of rotifer group Monogononta (sexual control for Bdelloidea) (Pejler, 1995) and ostracod group Cypridoidea (sexual control for Darwinulidae) (Martens et al., 2008) are planktonic; one of the two sexual lineages in genus *Lasaea* has planktonic larvae (Ó Foighil, 1988). Life in benthos could theoretically hamper and reduce the spread of parasites. Sessile organisms can resist hypothetical lethal epidemics postulated by the viral theory of background extinction more efficiently (Emiliani, 1993a, b).

On the other hand, paleontological studies show that species with planktonic larvae have decreased extinction rates in background extinction, particularly that species without planktonic larvae go extinct more rapidly (Jablonski, 1986). In *Lasaea*, it seems that lineages with direct development have a lower risk of extinction. These representatives also have a high chance of speciation after migration into the new area (Ó Foighil & Eernisse, 1988), but the main reason is probably that the direct developing lineages are surprisingly much better colonizers of distant areas thanks to the rafting on vegetation (Ó Foighil, 1989).

In a similar way to resting stages, the distribution of benthic or sedentary lifestyle among well supported AAs looks rather scarce on the first sight (3/8 cases). However, these 3 groups comprise all studied AAs that are (at least partially) associated with aquatic habitats. It is interesting that numerous contested aquatic AAs are also exclusively benthic: flatworm *Schmidtea polychroa* (Schmidt) (Pongratz et al., 2003), New Zealand mudsnail *Potamopyrgus antipodarum* (Gray) (Neiman et al., 2005), and ostracods *Heterocypris incongruens* (Ramdohr) and *Eucypris virens* (Jurine) (Butlin et al., 1998b; Martens, 1998).

### Life in the soil

Another adaptation widely distributed among AA groups is the inhabitancy of soil, especially deeper parts of the soil horizon. This tendency can be seen mainly in the AA mites from groups Oribatidae, Endeostigmata, and Trombidiformes (Karasawa & Hijii, 2008; Maraun et al., 2009; Walter, 2009). Bdelloidea and Darwinuloidea tend to be associated with semi-terrestrial habitats (Schön et al., 2009b). Moreover, AA Bdelloidea dominate among the soil rotifers (Pejler, 1995). Most representatives of Darwinulidae inhabit soil (respectively interstitial) too (Schön et al., 2009b). Taken together, 5/8 studied AA groups have numerous soil-inhabiting representatives and a tendency to inhabit soil. Moreover, Bdelloidea dominate there over its sexual control and AA mites have a stronger tendency to inhabit soil (Oribatidae) or its deep horizons (Endeostigmata, Trombidiformes) in comparison with their sexual controls.

Living in soil could, in a similar way to life in benthos, reduce the capacity of parasites to spread *(sensu* Emiliani, 1993a, b) Moreover, the tortuous character of the soil reduces any interactions of soil organisms and thus negatively affect not only parasitization but also predation and competition (Elliott et al., 1980; Murphy & Tate, 1996; Drake et al., 1998; Fisher et al., 2003; Lavelle & Spain, 2003; Paul, 2007). Besides, soil is an abiotically very stable environment shielding its inhabitants from fluctuations in temperature and humidity, as well as from UV radiation, and could be very favourable for asexuals also for this reason (Pilato, 1979; Krivolutsky & Druk, 1986; Siepel, 1994). However, other explanations have also been proposed for the asexuals’ association with soil habitats. Oribatidae could suffer less intense selective pressures in the soil than in the arboreal environment where they have to respond to the co-evolving lichens, their main food source (Maraun et al., 2009). Asexuality can be also more advantageous in soil because of the difficulties with seeking out sexual partners, less effective pheromone dispersal etc. (Karasawa & Hijii, 2008). Numerous contested AAs are soil inhabitants too: Glomales (Croll & Sanders, 2009), tardigrades (Pilato, 1979; Jorgensen, Mobjerg & Kristensen, 2007), and *Meloidogyne* (Castagnonesereno et al., 1993).

Thus, for the reasons mentioned above, life in soil habitats probably erases most of the evolutionary advantages of sexuality. Selection pressures of biotic and abiotic environments are weak in soil: The abiotic environment is relatively more stable and homogeneous; the interactions with competitors, parasites, and predators are very limited both in intensity and frequency due to the tortuous and multidimensional character of the environment (the shortest way from point A to point B in soil is only rarely a straight line). The evolutionary consequences of living in the soil are similar to those of having the capacity to form durable persistent stages (see above). This is in accordance with the presence of noticeable evolutionary stasis of soil-inhabiting groups described by Pilato (1979). It can also be seen in Bdelloidea (Ricci, 1987; Poinar & Ricci, 1992) and Darwinulidae (Martens et al., 1998; Schön et al., 1998, 2009b). On the other hand, the main factor causing evolutionary stasis in these two groups may be the subjective homogeneity of the environment caused by the presence of dormant stages. However, evolutionary stasis can be also found in other soil-inhabiting AAs: Oribatidae (Krivolutsky & Druk, 1986; Norton, 1994; Heethoff et al., 2007) and other AA mites (Norton et al., 1993; Walter, 2009; Walter et al., 2009); globally in 5/8 studied AA groups. Taking into account the contested AA groups, it can be found in Glomales (Remy et al., 1994; Redecker, Kodner & Graham, 2000) and tardigrades (Pilato, 1979; Jorgensen et al., 2007). All of these groups are very uniform and have undergone almost no change in tens to hundreds of millions of years (Pilato, 1979).

### Absence of life strategies with intensive biotic interactions

It is noticeable that there are practically no typical predators and parasites among the AAs we studied—this property is characteristic for all 8 studied groups. Remarkably often they feed on dead organic matter or are autotrophic; parasites are almost absent, and in the case of a predatory lifestyle, they are phytophagous or filtering (see Table S3). One possible explanation is that they are unable to keep up in the co-evolutionary race with their sexual hosts or prey. Thus, they can be successful in the long term, especially in the case of a predatory lifestyle, only if they adopt (or are preadapted to) such nonspecific ecological strategies.

### Succumbing to domestication and delegation of concern for its own benefit to another biological entity

The tendency for asexual reproduction is particularly interesting in the contested AA fungi domesticated by ants (Formicidae) and bark beetles (Scolytinae). The ant symbionts are from the basidiomycete groups Tricholomataceae and Lepiotaceae (Mueller et al., 1998), whereas bark beetles domesticate the ambrosia fungi of the ascomycete group Ophiostomatales (Farrell et al., 2001). The association is particularly close in the ants. They care for the fungi intensively, remove fungal predators and parasites, and the founding queen always carries filamentous bacteria, which synthetize an antidote against the main fungal pathogen— ascomycete *Escovopsis* (Currie et al., 1999a, b) —not to mention the stable temperature and humidity in the nest. By doing so, they provide a very favourable, biotically and abiotically stable environment. Moreover, there is some evidence that they prevent fungi from their already minimal attempts at sexual reproduction. On the other hand, the situation may be more complicated because some of these fungi create sexual structures predominantly in the presence of ants (Mueller, 2002). This phenomenon provides an alternative view on some aspects of human agriculture. A lot of crops raised by humans are asexual or self-pollinating too, which probably facilitates their breeding but increases their susceptibility to parasites and pathogens (Flegr, 2002). Life in association with another organism that takes care of the symbiont can also be found in the contested AA group Glomales (Croll & Sanders, 2009) and various prokaryotic and eukaryotic endosymbionts, see e.g. Douglas (2010). However, it has not been found in any of the 8 well supported AA groups we studied, and its effect on the long-term maintenance of asexual reproduction thus remains only speculative.

## Conclusions

The analytical part of this study, i.e. the comparative analysis of the environment of AAs and their sexual relatives, supported the hypothesis that AA groups are associated with overall (biotically and abiotically) more homogeneous environments in comparison with their sister or closely related ecologically comparable clades. It consequently supported the theoretical concepts that postulate the essential advantage of sexual species in heterogeneous environments and consider the (biotic and abiotic, temporal and spatial) heterogeneity of the environment affecting the organisms to be one factor that can exhibit itself in many ways (Williams, 1975, pp. 145–146, 149–154, 169; Roughgarden, 1991; Flegr, 2010, 2013). Particular ecological adaptations, from which durable resting stages, life in the absence of intense biotic interactions, and the association with soil and benthic habitats are most notable, could be considered special cases of the general AAs’ association with overall homogeneous environments.

Therefore, the general notion that proposed theories of sexual reproduction (see Introduction) need not exclude each other, that the effects proposed by some or all of them might intertwine and affect individuals and evolutionary lineages simultaneously, or that they even may, ultimately, represent only different aspects of one more general explanation, seems to be well supported. Moreover, overall environmental heterogeneity, regardless of its complicated conceptualization and study, seems to be a suitable candidate for this hypothetical general explanation.

Most putative AA lineages are still critically understudied. One way of elaborating the foundations laid out by this study would be comparing the heterogeneity of environments in a broader spectrum of AA lineages as soon as more lineages are discovered or confirmed (e.g. the protist lineages proposed by Speijer et al., 2015). It would also be very desirable to investigate the ecology of *Lasaea, Timema*, and Darwinulidae in greater detail. Additionally, it would be appropriate to focus on the interaction of biotic and abiotic environmental heterogeneities and their effect on organisms. According to Flegr (2008, 2010, 2013), sexual groups should exhibit more pronounced evolutionary conservation of niches in comparison with asexuals—on the whole, they are expected to stick closely around the phenotype of their common ancestor. This hypothesis could be tested by comparing the variance of properties of individual species within an AA and its related sexual clade. It would be also possible to test whether the sexual species are able to survive under a wider range of conditions of the heterogeneous environment due to their high genetic variability and hypothetical “elastic” reaction on selection, as was suggested by Flegr (2008, 2010, 2013).

## Acknowledgments

We are grateful to Radka Symonova, Karel Janko, Miloslav Devetter, Russell Shiel, Roy Norton, Jaroslav Smrz, Jan Mourek, Lubomir Kovac, Vladimir Sustr, Peter Luptacik, Josef Stary, Miroslav Kolarik, Lukas Kratochvil, Oldrich Fatka, Miroslav Kovarik, Adam Petrusek, Ivan Cepicka, Vojtech Hampl and Marek Elias for their help with collecting information about particular asexual taxa and useful insights into the biology of these species. We would also like to thank Karel Kotrly, Vlasta Pachtova and Vojtech Zarsky for their help with acquiring some hardly accessible literary sources and Eva Priplatova, Lenka Priplatova, Jinka Bousova, Julie Nokola Novakova and Charlie Lotterman for the final revisions of our text.

## Ethics approval and consent to participate

Not applicable

## Consent for publication

Not applicable

## Availability of data and materials

All data generated or analysed during this study are included in this published article and its supporting information files.

## Authors′ contributions

JT gathered the data, prepared the figures and tables and was the greatest contributor in writing the manuscript. JF contributed the analysis tools. Both JT and JF conceived and designed the study, analysed the data and reviewed the drafts of the paper. All authors read and approved the final manuscript.

## Additional files

**Additional file 1**: E&E supporting information.pdf (TITLE: 3 Supplemental tables and Supplemental review of AA ecology. DESRCRIPTION: Three Supplemental tables mentioned in the manuscript and Supplemental review that corroborates the claims in Supplemental table 3 and explains them in more detail).

## Footnotes

1 High genetic diversity of Bdelloidea in gene coxl is not very surprising in the light of severe DNA breaks that originate during anhydrobiosis, following repairs of these breaks and consequent intensive horizontal gene transfer (see e.g. Gladyshev et al., 2008).

## References

Becerra M, Brichette I, Garcia C. 1999. Short-term evolution of competition between genetically homogeneous and heterogeneous populations of Drosophila melanogaster. Evolutionary Ecology Research 1(5): 567–579.

Becks L, Agrawal AF. 2010. Higher rates of sex evolve in spatially heterogeneous environments. Nature 468(7320): 89–93.

Bell G. 1982. The masterpiece of nature: the evolution and genetics of sexuality. London, UK: Croom Helm.

Bell G. 1985. Two theories of sex and variation. Experientia 41(10): 1235–1245.

Bernstein H, Bernstein C. 2013. Evolutionary origin and adaptive function of meiosis. In: Bernstein H, Bernstein C, eds. Meiosis. InTech, 41–75. Available at: http://www.intechopen.com/books/meiosis/evolutionary-origin-and-adaptive-function-of-meiosis (accessed 23 may 2017). DOI: 10.5772/56557

Bettarel Y, Bouvy M, Dumont C, Sime-Ngando T. 2006. Virus-bacterium interactions in water and sediment of West African inland aquatic systems. Applied and Environmental Microbiology 72(8): 5274–5282.

Birky CJ. 2009. Sex and evolution in eukaryotes. Encyclopedia of Life Support Systems (EOLSS). Oxford, UK: Eolss Publishers. Available at: http://www.eolss.net/sample-chapters/c03/e6-183-14-00.pdf (accessed 23 may 2017).

Bogart JP, Bi K, Fu JZ, Noble DWA, Niedzwiecki J. 2007. Unisexual salamanders (genus Ambystoma) present a new reproductive mode for eukaryotes. Genome 50(2): 119–136.

Boschetti C, Pouchkina-Stantcheva N, Hoffmann P, Tunnacliffe A. 2011. Foreign genes and novel hydrophilic protein genes participate in the desiccation response of the bdelloid rotifer *Adineta ricciae*. Journal of Experimental Biology 214(1): 59–68.

Bratbak G, Egge JK, Heldal M. 1993. Viral mortality of the marine alga *Emiliania huxleyi* (Haptophyceae) and termination of algal blooms. Marine Ecology Progress Series 93: 39–48.

Bruvo R, Adolfsson S, Symonova R, Lamatsch D, Schön I, Jokela J, Butlin R, Muller S. 2011. Few parasites, and no evidence for *Wolbachia* infections, in a freshwater ostracod inhabiting temporary ponds. Biological Journal of the Linnean Society 102(1): 208–216.

Buckwold VE. 1994. Viral-induced extinctions unlikely. Nature 368(6471): 562.

Burt A, Bell G. 1987. Mammalian chiasma frequencies as a test of two theories of recombination. Nature 326(6115): 803–805.

Butlin R. 2002. The costs and benefits of sex: new insights from old asexual lineages. Nature Reviews Genetics 3(4): 311–317.

Butlin R, Schön I, Griffiths H. 1998a. Introduction to reproductive modes. In: Martens K, ed. Sex and Parthenogenesis: Evolutionary Ecology of Reproductive Modes in Non-Marine Ostracods. Leiden: Backhuys, 1–24.

Butlin R, Schön I, Martens K. 1998b. Asexual reproduction in nonmarine ostracods. Heredity 81(5): 473–480.

Butlin R, Schön I, Martens K. 1999. Origin, age and diversity of clones. Journal of Evolutionary Biology 12(6): 1020–1022.

Carbonel P, Colin J, Danielopol D, Loffler H, Neustrueva I. 1988. Paleoecology of limnic ostracodes: a review of some major topics. Palaeogeography Palaeoclimatology Palaeoecology 62(1–4): 413–461.

Castagnonesereno P, Piotte C, Uijthof J, Abad P, Wajnberg E, Vanlerberghemasutti F, Bongiovanni M, Dalmasso A. 1993. Phylogenetic relationships between amphimictic and parthenogenetic nematodes of the genus *Meloidogyne* as inferred from repetitive DNA analysis. Heredity 70(2): 195–204.

Colegrave N, Kaltz O, Bell G. 2002. The ecology and genetics of fitness in *Chlamydomonas*. VIII. The dynamics of adaptation to novel environments after a single episode of sex. Evolution 56(1): 14–21.

Coleman DA, Crossley DA, Hendrix PF. 2004. Fundamentals of Soil Ecology, Second edition. USA: Elsevier Academic Press.

Croll D, Sanders I. 2009. Recombination in *Glomus intraradices*, a supposed ancient asexual arbuscular mycorrhizal fungus. Bmc Evolutionary Biology 9(1).

Crow J. 1970. Genetic loads and the cost of natural selection. In: Kojima K, ed. Biomathematics. Volume 1. Mathematical topics in population genetics. Berlin: Springer-Verlag, 128–177.

Currie C, Mueller U, Malloch D. 1999a. The agricultural pathology of ant fungus gardens. Proceedings of the National Academy of Sciences of the United States of America 96(14): 7998–8002.

Currie C, Scott J, Summerbell R, Malloch D. 1999b. Fungus-growing ants use antibiotic-producing bacteria to control garden parasites. Nature 398(6729): 701–704.

Danovaro R, Dell’Anno A, Corinaldesi C, Magagnini M, Noble R, Tamburini C, Weinbauer M. 2008. Major viral impact on the functioning of benthic deep-sea ecosystems. Nature 454(7208): 1084–1087.

Dawkins R, Krebs JR. 1979. Arms races between and within species. Proceedings of the Royal Society B-Biological Sciences 205(1161): 489–511.

Debortoli N, Li X, Eyres I, Fontaneto D, Hespeels B, Tang CQ, Flot J, Van Doninck K. 2016. Genetic exchange among bdelloid rotifers is more likely due to horizontal gene transfer than to meiotic sex. Current Biology 26(6): 723–732.

de Castro F, Bolker B. 2005. Mechanisms of disease-induced extinction. Ecology Letters 8(1): 117–126.

de Meeus T, Prugnolle F, Agnew P. 2007. Asexual reproduction: genetics and evolutionary aspects. Cellular and Molecular Life Sciences 64(11): 1355–1372.

Devetter M, Scholl K. 2014. Hydrobiont animals in floodplain soil: are they positively or negatively affected by flooding? Soil Biology & Biochemistry 69: 393–397.

Dole-Olivier M, Galassi D, Marmonier P, Des Chatelliers M. 2000. The biology and ecology of lotic microcrustaceans. Freshwater Biology 44(1): 63–91.

Domes K, Scheu S, Maraun M. 2007. Resources and sex: soil re-colonization by sexual and parthenogenetic oribatid mites. Pedobiologia 51(1): 1–11.

Douglas A. 2010. The Symbiotic Habit. Princeton, NJ: Princeton University Press.

Drake LA, Choi KH, Haskell AE, Dobbs FC. 1998. Vertical profiles of virus-like particles and bacteria in the water column and sediments of Chesapeake Bay, USA. Aquatic Microbial Ecology 16(1): 17–25.

Elliott ET, Anderson RV, Coleman DC, Cole CV. 1980. Habitable pore space and microbial trophic interactions. Oikos 35(3): 327–335.

Emiliani C. 1982. Extinctive evolution: extinctive and competitive evolution combine into a unified model of evolution. Journal of Teoretical Biology 97(1): 13–33.

Emiliani C. 1993a. Extinction and viruses. Biosystems 31(2–3): 155–159.

Emiliani C. 1993b. Viral extinctions in deep-sea species. Nature 366(6452): 217–218.

Etter RJ, Rex MA, Chase MR, Quattro JM. 2005. Population differentiation decreases with depth in deep-sea bivalves. Evolution 59(7): 1479–1491.

Farrar D. 1978. Problems in the identity and origin of the Appalachian *Vittaria* gametophyte, a sporophyteless fern of the eastern United States. American Journal of Botany 65(1): 1–12.

Farrar D. 1990. Species and evolution in asexually reproducing independent fern gametophytes. Systematic Botany 15(1): 98–111.

Farrar D. 1992. *Trichomanes intricatum:* the independent *Trichomanes* gametophyte in the eastern United States. American Fern Journal 82(2): 68–74.

Farrar D. 1998. The tropical flora of rockhouse cliff formations in the eastern United States. Journal of the Torrey Botanical Society 125(2): 91–108.

Farrell BD, Sequeira AS, O’Meara BC, Normark BB, Chung JH, Jordal BH. 2001. The evolution of agriculture in beetles (Curculionidae: Scolytinae and Platypodinae). Evolution 55(10): 2011–2027.

Filippini M, Buesing N, Bettarel Y, Sime-Ngando T, Gessner MO. 2006. Infection paradox: high abundance but low impact of freshwater benthic viruses. Applied and Environmental Microbiology 72(7): 4893–4898.

Fischer O & Schmid-Hempel P. 2005. Selection by parasites may increase host recombination frequency. Biology Letters 1(2): 193–195.

Fisher R. 2003. The Genetical Theory of Natural Selection: A Complete Variorum Edition. New York, NY: Oxford University Press.

Fisher UR, Wieltschnig C, Kirschner AKT, Velimirov B. 2003. Does virus-induced lysis contribute significantly to bacterial mortality in the oxygenated sediment layer of shallow oxbow lakes? Applied and Environmental Microbiology 69(9): 5281–5289.

Flegr J. 2002. Was Lysenko (partly) right? Michurinist biology in the view of modern plant physiology and genetics. Rivista Di Biologia-Biology Forum 95: 259–271.

Flegr J. 2008. Frozen evolution: Or, that’s not the way it is, mr. Darwin - Farewell to selfish gene. USA: Createspace Independent Pub.

Flegr J. 2010. Elastic, not plastic species: frozen plasticity theory and the origin of adaptive evolution in sexually reproducing organisms. Biology Direct 5(1).

Flegr J. 2013. Microevolutionary, macroevolutionary, ecological and taxonomical implications of punctuational theories of adaptive evolution. Biology Direct 8(1).

Fuhrman JA. 1999. Marine viruses and their biogeochemical and ecological effects. Nature 399(6736): 541–548

Garcia C, Toro M. 1992. Sib competition in *Tribolium:* a test of the elbow-room model. Heredity 68(6): 529–536.

Gladyshev E, Meselson M. 2008. Extreme resistance of bdelloid rotifers to ionizing radiation. Proceedings of the National Academy of Sciences of the United States of America 105(13): 5139–5144.

Glesener R, Tilman D. 1978. Sexuality and the components of environmental uncertainty: clues from geographic parthenogenesis in terrestrial animals. The American Naturalist 112(986): 659–673.

Gorelick R, Carpinone J. 2009. Origin and maintenance of sex: the evolutionary joys of self sex. Biological Journal of the Linnean Society 98(4): 707–728.

Griffiths H, Butlin R. 1995. A timescale for sex versus parthenogenesis: evidence from subfossil ostracods. Proceedings of the Royal Society of London Series B-Biological Sciences 260(1357): 65–71.

Hamilton W, Axelrod R, Tanese R. 1990. Sexual reproduction as an adaptation to resist parasites (a review). Proceedings of the National Academy of Sciences of the United States of America 87(9): 3566–3573.

Heethoff M, Domes K, Laumann M, Maraun M, Norton RA, Scheu S. 2007. High genetic divergences indicate ancient separation of parthenogenetic lineages of the oribatid mite *Platynothrus peltifer* (Acari, Oribatida). Journal of Evolutionary Biology 20(1): 392–402.

Hörandl E. 2006. The complex causality of geographical parthenogenesis. New Phytologist 171(3): 525–538.

Hörandl E. 2009. Geographical parthenogenesis: opportunities for asexuality. In: Schön I, Martens K, van Dijk P, eds. Lost Sex: The Evolutionary Biology of Parthenogenesis. Dordrecht: Springer Science+Business Media B.V., 161–186.

Jablonski D. 1986. Background and mass extinctions: the alternation of macroevolutionary regimes. Science 231(4734): 129–133.

Janko K, Drozd P, Eisner J. 2011. Do clones degenerate over time? Explaining the genetic variability of asexuals through population genetic models. Biology Direct 6(1).

Janko K, Drozd P, Flegr J, Pannell J. 2008. Clonal turnover versus clonal decay: a null model for observed patterns of asexual longevity, diversity and distribution. Evolution 62(5): 1264–1270.

Jorgensen A, Mobjerg N, Kristensen R. 2007. Molecular study of the tardigrade *Echiniscus testudo* (Echiniscidae) reveals low DNA sequence diversity over a large geographical area. Journal of Limnology 66(1s): 77–83.

Judson OP, Normark BB. 1996. Ancient asexual scandals. Trends in Ecology & Evolution 11(2): 41–46.

Kaltz O, Bell G. 2002. The ecology and genetics of fitness in *Chlamydomonas.* XII. Repeated sexual episodes increase rates of adaptation to novel environments. Evolution 56(9): 1743–1753.

Karasawa S, Hijii N. 2008. Vertical stratification of oribatid (Acari: Oribatida) communities in relation to their morphological and life-history traits and tree structures in a subtropical forest in Japan. Ecological Research 23(1): 57–69.

Koella J. 1993. Ecological correlates of chiasma frequency and recombination index of plants. Biological Journal of the Linnean Society 48(3): 227–238.

Kondrashov A. 1982. Selection against harmful mutations in large sexual and asexual populations. Genetical Research 40(03): 325–332.

Kondrashov A. 1993. Classification of hypotheses on the advantage of amphimixis. Journal of Heredity 84(5): 372–387.

Krivolutsky D, Druk A. 1986. Fossil oribatid mites. Annual Review of Entomology 31(1): 533–545.

Lavelle P, Spain AV. 2003. Soil Ecology. New York, USA: Kluwer Academic Publishers.

Law J, Crespi B. 2002. Recent and ancient asexuality in *Timema* walkingsticks. Evolution 56(8): 1711–1717.

Lehtonen J, Jennions M, Kokko H. 2012. The many costs of sex. Trends in Ecology & Evolution 27(3): 172–178.

Lewis J, Wolpert L. 1979. Diploidy, evolution and sex. Journal of Theoretical Biology 78(3): 425–438.

Li H, Reynolds J. 1995. On definition and quantification of heterogeneity. Oikos 73(2): 280–284.

Mantovani B, Passamonti M, Scali V. 2001. The mitochondrial cytochrome oxidase II gene in *Bacillus* stick insects: ancestry of hybrids, androgenesis, and phylogenetic relationships. Molecular Phylogenetics and Evolution 19(1): 157–163.

Maraun M, Erdmann G, Schulz G, Norton RA, Scheu S, Domes K. 2009. Multiple convergent evolution of arboreal life in oribatid mites indicates the primacy of ecology. Proceedings of the Royal Society B-Biological Sciences 276(1671): 3219–3227.

Maraun M, Norton RA, Ehnes RB, Scheu S, Erdmann G. 2012. Positive correlation between density and parthenogenetic reproduction in oribatid mites (Acari) supports the structured resource theory of sexual reproduction. Evolutionary Ecology Research 14(3): 311–323.

Martens K. 1998. Sex and ostracods: a new synthesis. In: Martens K, ed. Sex and Parthenogenesis: Evolutionary Ecology of Reproductive Modes in Non-Marine Ostracds. Leiden: Backhuys, 295–321.

Martens K, Horne D, Griffiths H. 1998. Age and diversity of non-marine ostracods. In: Martens K, ed. Sex and Parthenogenesis: Evolutionary Ecology of Reproductive Modes in NonMarine Ostracds. Leiden: Backhuys, 37–55.

Martens K, Schön I. 2000. The importance of habitat stability for the prevalence of sexual reproduction. In: Minoura K, ed. Lake Baikal: a Mirror in Time and Space for Understanding Global Change Processes. Yokohama Symposium 1998. Amsterdam: Elsevier, 324–330.

Martens K, Schön I, Meisch C, Horne D. 2008. Global diversity of ostracods (Ostracoda, Crustacea) in freshwater. Hydrobiologia 595(1): 185–193.

Maynard Smith J. 1978. The evolution of sex. Cambridge, UK: Cambridge University Press.

Maynard Smith J. 1993. The theory of evolution. Cambridge, UK: Cambridge University Press.

McDonald JH. 2014. Handbook of Biological Statistics (3rd ed.). Baltimore, USA: Sparky House Publishing.

Meirmans S, Strand R. 2010. Why are there so many theories for sex, and what do we do with them? Journal of Heredity 101(1s): S3–S12.

Mikheyev AS, Mueller UG, Abbot P. 2006. Cryptic sex and many-to-one colevolution in the fungus-growing ant symbiosis. Proceedings of the National Academy of Sciences of the United States of America 103(28): 10702–10706.

Mobjerg N, Halberg K, Jorgensen A, Persson D, Bjorn M, Ramlov H, Kristensen R. 2011. Survival in extreme environments – on the current knowledge of adaptations in tardigrades. Acta Physiologica 202(3): 409–420.

Morton JE, Boney AD, Corner EDS. 1957. The adaptations of *Lasaea rubra* (Montagu), a small intertidal lamellibranch. Journal of the Marine Biological Association of the United Kingdom 36(02): 383–405.

Mueller UG. 2002. Ant versus fungus versus mutualism: ant–cultivar conflict and the deconstruction of the attine ant–fungus symbiosis. American Naturalist 160(4s): S67–S98.

Mueller UG, Rehner SA, Schultz TR. 1998. The evolution of agriculture in ants. Science 281(5385): 2034–2038.

Muller H. 1932. Some genetic aspects of sex. The American Naturalist 66(703): 118–138.

Muller H. 1964. The relation of recombination to mutational advance. Mutation Research 1(1): 2–9.

Murphy SL, Tate RL. 1996. Bacterial movement through soil. In: Stozsky G & Bollag JM, ed. Soil Biochemistry Vol. 9. New York: Marcel Dekker, 253–286.

Neiman M, Jokela J, Lively C. 2005. Variation in asexual lineage age in *Potamopyrgus antipodarum*, a New Zealand snail. Evolution 59(9): 1945–1952.

Neiman M, Koskella B. 2009. Sex and the Red queen. In: Schön I, Martens K, van Dijk P, eds. Lost sex: the Evolutionary Biology of Parthenogenesis. Dordrecht: Springer Science+Business Media B.V., 133–159.

Neiman M, Meirmans S, Meirmans P, Schlichting C, Mousseau T. 2009. What can asexual lineage age tell us about the maintenance of sex? Annals of the New York Academy of Sciences 1168(1): 185–200.

Neiman M, Schwander T. 2011. Using parthenogenetic lineages to identify advantages of sex. Evolutionary Biology 38(2): 115–123.

Normark B, Judson O, Moran N. 2003. Genomic signatures of ancient asexual lineages. Biological Journal of the Linnean Society 79(1): 69–84.

Norton R. 1994. Evolutionary aspects of oribatid mite life histories and consequences for the origin of the Astigmata. In: Houck M, ed. Mites. Ecological and Evolutionary Analyses of Life-History Patterns. New York: Chapman & Hall, 99–135.

Norton R, Kethley J, Johnston D, O’Connor B. 1993. Phylogenetic perspectives on genetic systems and reproductive modes of mites. In: Wrensch D, Ebbert M, eds. Evolution and diversity of sex ratio in insects and mites. New York: Chapman and Hall, 8–99.

Nunney L. 1989. The maintenance of sex by group selection. Evolution 43(2): 245–257.

Otto S, Lenormand T. 2002. Resolving the paradox of sex and recombination. Nature Reviews Genetics 3(4): 252–261.

Ó Foighil D. 1988. Random mating and planktotrophic larval development in the brooding hermaphroditic clam *Lasaea australis* (Lamarck, 1818). Veliger 31(3–4): 214–221.

Ó Foighil D. 1989. Planktotrophic larval development is associated with a restricted geographic range in *Lasaea*, a genus of brooding, hermaphroditic bivalves. Marine Biology 103(3): 349–358.

Ó Foighil D, Eernisse D. 1988. Geographically widespread, non-hybridizing, sympatric strains of the hermaphroditic, brooding clam *Lasaea* in the northeastern Pacific Ocean. Biological Bulletin 175(2): 218–229.

Paul EA. 2007. Soil Microbiology, Ecology and Biochemistry. USA: Academic Press.

Pejler B. 1995. Relation to habitat in rotifers. Hydrobiologia 313(1): 267–278.

Pilato G. 1979. Correlations between cryptobiosis and other biological characteristics in some soil animals. Italian Journal of Zoology 46(4): 319–332.

Poinar GO, Ricci C. 1992. Bdelloid rotifers in Dominican amber: evidence for parthenogenetic continuity. Experientia 48(4): 408–410.

Pongratz N, Storhas M, Carranza S, Michiels N. 2003. Phylogeography of competing sexual and parthenogenetic forms of a freshwater flatworm: patterns and explanations. Bmc Evolutionary Biology 3(1).

Pontecorvo D. 1954. Mitotic recombination in the genetic systems of filamentous fungi. Caryologia 6: 192–200.

Quesada CA, Miranda AC, Hodnett MG, Santos AJB, Miranda HS, Breyer LM. 2004. Seasonal and depth variation of soil moisture in a burned open savanna (campo sujo) in central Brazil. Ecological Applications 14(4s): 33–41.

Redecker D, Kodner R, Graham L. 2000. Glomalean fungi from the Ordovician. Science 289(5486): 1920–1921.

Redfield R. 2001. Do bacteria have sex? Nature Reviews Genetics 2(8): 634–639.

Remy W, Taylor T, Hass H, Kerp H. 1994. Four hundred-million-year-old vesicular arbuscular mycorrhizae. Proceedings of the National Academy of Sciences of the United States of America 91(25): 11841–11843.

Renaut S, Replansky T, Heppleston A, Bell G. 2006. The ecology and genetics of fitness in *Chlamydomonas.* XIII. The fitness of long-term sexual and asexual populations in benign environments. Evolution 60(11): 2272–2279.

Ricci C. 2001. Dormancy patterns in rotifers. Hydrobiologia 446(1): 1–11.

Ricci C, Balsamo M. 2000. The biology and ecology of lotic rotifers and gastrotrichs. Freshwater Biology 44(1): 15–28.

Ricci CN. 1987. Ecology of bdelloids: how to be successful. Hydrobiologia 147(1): 117–127.

Robeson MS, King AJ, Freeman KR, Birky CW, Martin AP, Schmidt SK. 2011. Soil rotifer communities are extremely diverse globally but spatially autocorrelated locally. Proceedings of the National Academy of Sciences of the United States of America 108(11): 4406–4410.

Rohde K. 1986. Differences in species diversity of monogenea between the pacific and atlantic oceans. Hydrobiologia 137(1): 21–28.

Rohde K, Heap M. 1998. Latitudinal differences in species and community richness and in community structure of metazoan endo- and ectoparasites of marine teleost fish. International Journal For Parasitology 28(3): 461–474.

Roughgarden J. 1991. The evolution of sex. American Naturalist 138(4): 934–953.

R_Core_Team. 2014. R: A language and environment for statistical computing, version 3.1.2. Vienna, Austria: R Foundation for Statistical Computing. Available at: https://www.r-project.org (accessed 23 may 2016).

Schurko A, Neiman M, Logsdon J. 2009. Signs of sex: what we know and how we know it. Trends in Ecology & Evolution 24(4): 208–217.

Schwander T, Crespi BJ. 2009. Twigs on the tree of life? Neutral and selective models for integrating macroevolutionary patterns with microevolutionary processes in the analysis of asexuality. Molecular Ecology 18(1): 28–42.

Schwander T, Henry L, Crespi B. 2011. Molecular evidence for ancient asexuality in *Timema* stick insects. Current Biology 21(13): 1129–1134.

Sheldon P. 1996. Plus ça change—a model for stasis and evolution in different environments. Palaeogeography, Palaeoclimatology, Palaeoecology 127(1–4): 209–227.

Schön I, Butlin R, Griffiths H, Martens K. 1998. Slow molecular evolution in an ancient asexual ostracod. Proceedings of the Royal Society of London Series B-Biological Sciences 265(1392): 235–242.

Schön I, Martens K. 2004. Adaptive, pre-adaptive and non-adaptive components of radiations in ancient lakes: a review. Organisms Diversity & Evolution 4(3): 137–156.

Schön I, Martens K, Rossi V. 1996. Ancient asexuals: scandal or artifact? Trends in Ecology and Evolution 11(7): 296–297.

Schön I, Martens K, van Dijk P. 2009a. Lost Sex: The Evolutionary Biology of Parthenogenesis. Dordrecht: Springer Science+Business Media B.V.

Schön I, Rossetti G, Martens K. 2009b. Darwinulid ostracods: ancient asexual scandals or scandalous gossip? In: Schon I, Martens K, van Dijk P, eds. Lost Sex: The Evolutionary Biology of Parthenogenesis: Dordrecht: Springer Science+Business Media B.V., 217–240.

Schwander T. 2016. Evolution: The end of an ancient asexual scandal. Current Biology 26(6): R233–R235.

Siepel H. 1994. Life-history tactics of soil microarthropods. Biology and Fertility of Soils 18(4): 263–278.

Siepel H. 1996. Biodiversity of soil microarthropods: the filtering of species. Biodiversity & Conservation 5(2): 251–260.

Smith J. 1980. Selection for recombination in a polygenic model. Genetical Research 35(03): 269–277.

Speijer D, Lukes J, Elias M. 2015. Sex is a ubiquitous, ancient, and inherent attribute of eukaryotic life. Proceedings of the National Academy of Sciences of the United States of America 112(29): 8827–8834.

Sterrer W. 2002. On the origin of sex as vaccination. Journal of Theoretical Biology 216(4): 387–396.

Suttle CS. 2005. Viruses in the sea. Nature 437(7057): 356–361.

Suttle CS. 2007. Marine viruses—major players in the global ecosystem. Nature Reviews Microbiology 5(10): 801–812.

Suttle CS, Chan AM, Cottrell MT. 1990. Infection of phytoplankton by viruses and reduction of primary productivity. Nature 347(6292): 467–469.

Thornhill R, Fincher C. 2013. The comparative method in cross-cultural and cross-species research. Evolutionary Biology 40(4): 480–493.

Tobler M, Schlupp I, de Leon F, Glaubrecht M, Plath M. 2007. Extreme habitats as refuge from parasite infections? Evidence from an extremophile fish. Acta Oecologica 31(3): 270–275.

Tokeshi M. 1999. Species Coexistence: Ecological and Evolutionary Perspectives. Oxford, UK: Blackwell Science Ltd.

Turgeon J, Hebert P. 1994. Evolutionary interactions between sexual and all-female taxa of *Cyprinotus* (Ostracoda: Cyprididae). Evolution 48(6): 1855–1865.

Van Dijk P. 2009. Apomixis: basics for non-botanists. In: Schön I, Martens K, van Dijk P, eds. Lost Sex: The Evolutionary Biology of Parthenogenesis. Dordrecht: Springer Science+Business Media B.V., 47–62.

van Raay T, Crease T. 1995. Mitochondrial DNA diversity in an apomictic *Daphnia complex* from the Canadian High Arctic. Molecular Ecology 4(2): 149–161.

Vanhaecke P, Siddall S, Sorgeloos P. 1984. International study on *Artemia.* XXXII. Combined effects of temperature and salinity on the survival of *Artemia* of various geographical origin. Journal of Experimental Marine Biology and Ecology 80(3): 259–275.

von Uexküll J. 1982. The theory of Meaning. Semiotica 42(1): 25–82.

Vrijenhoek R. 1984. The evolution of clonal diversity in *Poeciliopsis.* In: Turner B, ed. Evolutionary Genetics of Fishes. New York: Plenum Press, 399–429.

Vrijenhoek R, Parker E. 2009. Geographical parthenogenesis: general purpose genotypes and frozen niche variation. In: Schön I, Martens K, van Dijk P, eds. Lost Sex: The Evolutionary Biology of Parthenogenesis: Dordrecht: Springer Science+Business Media B.V., 99–131.

Walter D. 2009. Suborder Endeostigmata. In: Krantz G, Walter D, eds. A manual of acarology. Lubbock: Texas Tech University Press, 421–429.

Walter D, Lindquist E, Smith I, Cook D, Krantz G. 2009. Order Trombidiformes. In: Krantz G, Walter D, eds. A manual of acarology. Lubbock: Texas Tech University Press, 233–420.

Wallwork, JA. 1970. Ecology of soil animals. London, UK: McGRAW-HILL Publishing Company Limited.

West S, Lively C, Read A. 1999. A pluralist approach to sex and recombination. Journal of Evolutionary Biology 12(6): 1003–1012.

Williams GC. 1975. Sex and evolution. Princeton, NJ: Princeton University Press.

Wilson CG. 2011. Desiccation-tolerance in bdelloid rotifers facilitates spatiotemporal escape from multiple species of parasitic fungi. Biological Journal of the Linnean Society 104(3): 564–574.

Wommack KE, Colwell RR. 2000. Virioplankton: Viruses in Aquatic Ecosystems. Microbiology and Molecular Biology Reviews 64(1): 69–114.

